# Atypical flexibility in dynamic functional connectivity quantifies the severity in autism spectrum disorder

**DOI:** 10.1101/387886

**Authors:** Vatika Harlalka, Raju S. Bapi, P.K. Vinod, Dipanjan Roy

## Abstract

Resting-state functional connectivity (FC) analyses have shown atypical connectivity in autism spectrum disorder (ASD) as compared to typically developing (TD). However, this view emerges from investigating static FC overlooking the age, disease phenotype and their interaction in the whole brain transient connectivity patterns. Contrasting with most extant literature in the present study, we investigated precisely how age and disease phenotypes factors into dynamic changes in functional connectivity of TD and ASD using resting-state functional magnetic resonance imaging (rs-fMRI) data stratified into three cohorts: children (7–11 years) and adolescents (12–17 years), and adults (18+) for the analysis. The dynamic variability in the connection strength and the modular organization in terms of measures: flexibility, cohesion strength and disjointness were explored for each subject to characterize the differences between ASD and TD.

In ASD, we observed significantly higher inter-subject dynamic variability in connection strength as compared to TD. This hypervariability relates to the symptom severity in ASD. We found that whole-brain flexibility correlates with static modularity only in TD. Further, we observed a core-periphery organization in the resting-state, with Sensorimotor and Visual regions in the rigid core; and DMN and attention areas in the flexible periphery. TD also develops a more cohesive organization of sensorimotor areas. However, in ASD we found a strong positive correlation of symptom severity with the flexibility of rigid areas and with disjointness of sensorimotor areas. The regions of the brain showing the high predictive power of symptom severity were distributed across the cortex, with stronger bearings in the frontal, motor and occipital cortices. Our study demonstrates that the dynamic framework best characterizes the variability in ASD.

## 1. Introduction

Autism spectrum disorder (ASD) is a lifelong neurodevelopmental disorder characterized by deficits in social communication skills, restricted and repetitive behaviour. It encompasses a range of disorders including Asperger’s and pervasive developmental disorder. Resting state functional MRI, which measures blood oxygen level-dependant signals (BOLD) (Deco et al., 2013; Lee et al., 2013) has been used to study both physiology and pathology. The statistical analysis of the matrix obtained by using Pearson cross-correlation of the regional BOLD time series has been found to reveal several properties that are of clinical relevance. Recently, a number of studies have explored brain networks as complex graphs (Sporns, 2013; van den Heuvel and Hulshoff Pol, 2010). Graph-theoretical analyses have shown that the functional connectivity in the human brain is divided into well-organized modules or subnetworks which are densely connected within themselves and sparsely connected to each other. Disease and age affect this modular organization (Chen et al., 2013; Song et al., 2014; Ye et al., 2015). Jie Song et al (2014) reported that modularity decreases with aging, suggesting less distinct functional divisions and specialization across whole brain networks. Moreover, they found a decline in cognitive functions but maintenance of primary information processing in normal healthy aging (Song et al., 2014).

Several studies on neurocognitive diseases have reported atypical fluctuation in modularity. Recently, in ASD, alterations in the network properties (global and local efficiency, assortativity, clustering coefficients, characteristic path length, small world properties etc.) with maturation and disease were found in our and previous studies (Harlalka et al., 2018; Henry et al., 2017; Rudie et al., 2013). Using static networks, a significant decrease in modularity has been observed. Both the functional connectivity *between* major networks (i.e. functional segregation) and connectivity *within* different networks (i.e. functional integration) are altered in ASD (Rudie et al., 2013, 2012). Studies have shown underconnectivity in various functional networks, especially the Default Mode Network (DMN) (Hahamy et al., 2015; Yerys et al., 2015).

However, in recent times, it has been found that static functional connectivity has several shortcomings. It does not sufficiently incorporate the time-varying (or dynamic) changes that occur through the brain scan (Chang and Glover, 2010). Dynamic functional connectivity has been shown to reveal patterns of brain states that occur commonly as well as the transitions between them (Damaraju et al., 2014; Preti et al., 2017). They also give an idea about dynamic reconfiguration that occurs during tasks (Bassett et al., 2011; Braun et al., 2016; Gerraty et al., 2018). A common approach used to perform dynamic connectivity analysis is to divide the scan session into overlapping sub-intervals or windows, and calculate a full correlation matrix for each sub-interval. The correlation matrix for each sub-interval is taken as an estimate of the instantaneous functional connectivity. This sliding window or tapered sliding window approach has been widely used to study dynamic connectivity of functional brain networks estimated from fMRI BOLD data (Park et al., 2017; Xu and Lindquist, 2015). With recent studies reporting that modularity of time-resolved functional networks varies on shorter timescales (Betzel et al., 2017, 2016), there is a possibility of tracking instantaneous changes in functional connectivity between brain regions. Particularly, while static communities represent sub-networks densely connected among themselves and sparsely connected to the rest of the brain, community structures in dynamic networks would project the ongoing changes in communities over time (Bassett et al., 2011; Cole et al., 2013).

Recent studies have explored the dynamic nature of atypical information processing in ASD (Chen et al., 2017; de Lacy et al., 2017; Falahpour et al., 2016; Rashid et al., 2018; Watanabe and Rees, 2017). Watanabe et al (2017) reported that ASD shows fewer neural transitions compared to those of neurotypical controls, and such atypically stable brain dynamics underlie both core symptoms in ASD and general cognitive ability (Watanabe and Rees, 2017). Some studies reported altered dynamic functional connectivity between specific areas (Falahpour et al., 2016). Most studies on whole-brain have found dominant brain states as well as their dwell times. ASD is shown to spend more time in globally disconnected states with less number of state transitions as compared to TD (de Lacy et al., 2017; Rashid et al., 2018).

Flexibility of brain also provides an alternate way of quantifying the dynamic changes in fMRI studies (Garcia et al., 2018). Intuitively, it can be thought of as a statistic to quantify the amount of reconfiguration in functional connectivity patterns that a brain region displays over time. It has been applied to obtain insights into altered dynamic pattern in various diseased states like schizophrenia (Braun et al., 2016) and epilepsy (Tailby et al., 2018). The influence of resting-state flexibility on disease and age has not been investigated, particularly in the setting of ASD.

Therefore, we adopt this framework to investigate the dynamic changes in functional activity of TD and ASD. We stratified the data (ASD and TD) into groups of children, adolescents and adults. We attempted to quantify the variability in dynamic FC in two plausible ways.

Firstly, we quantified the variability in the connection strength. Secondly, dynamic metrics, flexibility, cohesion and disjointness, were used to study the differences between ASD and TD. The pipeline is summarized in **Fig 1.**

**Fig 1.**
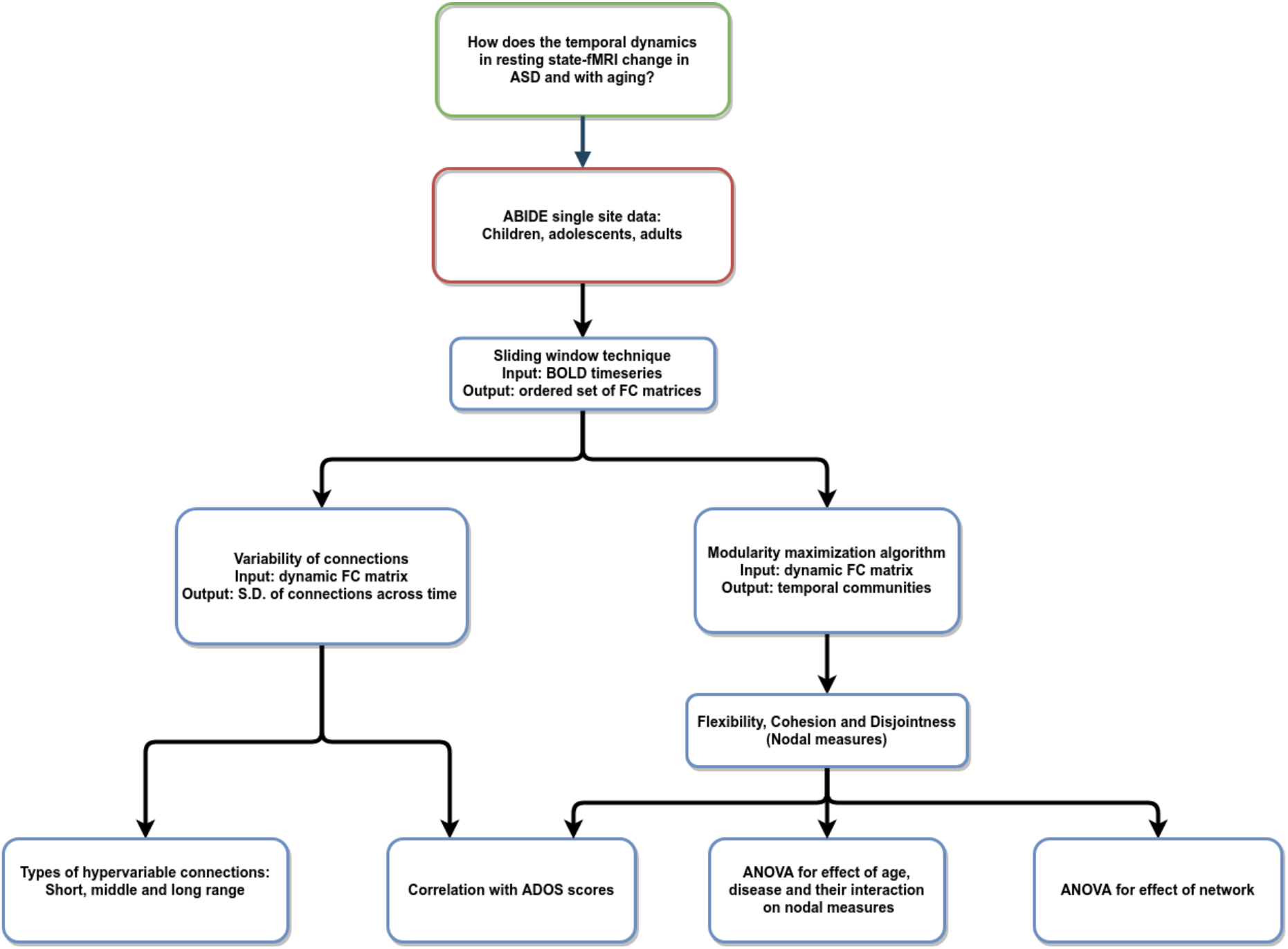
Pipeline of the analysis: resting-state fMRI data used to analyze varying temporal dynamics in ASD and TD.

Our study shows that ASD follows a very different dynamic trajectory as compared to TD. In ASD we observed significantly higher inter-subject dynamic variability in connection strength as compared to TD. This hypervariability relates to the symptom severity in ASD. We found that whole-brain flexibility correlates with static modularity only in TD. We observed a core-periphery organization in the resting-state, with Sensorimotor and Visual regions in the rigid core; and DMN and attention areas in the flexible periphery. TD also develops a more cohesive organization of sensorimotor areas. However, in ASD we found a strong positive correlation of symptom severity with flexibility in rigid areas and with disjointness of sensorimotor areas. The regions of the brain showing high predictive power of symptom severity were distributed across the cortex, with stronger bearings in the frontal, motor and occipital cortices.

## 2. Materials and methods

### 2.1 Data Acquisition and Preprocessing

We used single-site data (NYU) from the ABIDE Preprocessed Initiative (Cameron et al., 2013). Institutional review board approval was provided by each data contributor in the ABIDE database. Detailed recruitment and assessment protocols and inclusion criteria are all available on the ABIDE website. The data was preprocessed using the DPARSF pipeline (Data Preprocessing Assistant for Resting State fMRI) (Yan, 2010). The first 10 images were discarded for steady-state longitudinal magnetization. Temporal and head motion correction was applied on the remaining images, which were then warped into standard Montreal Neurological Institute (MNI) space. All normalized images were smoothed with a Gaussian kernel (full width at half maximum 5 mm) and then detrended. Signals from white matter, cerebrospinal fluid, and 24 rigid body motion parameters were regressed out of the data. The white matter and cerebrospinal fluid (CSF) masks were obtained from the tissue probability maps. We did not apply global signal regression. Finally a bandpass filter (0.01–0.1 Hz) was applied on the regressed time series (Liu and Duyn, 2013). Details of the preprocessing steps can be found here:http://preprocessed-connectomes-project.org/abide/dparsf.html.

Participants data was visually inspected by 3 functional raters. Subjects were excluded if (1) Any of the 3 raters gave it a ‘Maybe’ or a ‘Fail’ rating. (2) failed visual inspection of anatomical images and surfaces; (2) mean framewise displacement > 0.1 mm. Further, the data was age-stratified into three cohorts: young children under 11 years of age (*n* = 52), adolescents from 11–18 years of age (*n* = 56), and adults over 18 years of age (*n* = 36) (**Table 1**). Overall, within each age group, we did not find significant differences in within-group properties for age and IQ between ASD and TD subjects. Non-parametric t-tests were used to calculate differences in mean relative motion and IQ scores. While motion was slightly higher in children compares to adolescents and adults, we found that there were no significant differences in these measures between ASD and TD.

**Table 1.** Demographic details. Details about demographics of the samples included in this study.

|  | ASD | TD | p value |
| --- | --- | --- | --- |
| Children |  |  |  |
| Age: mean (SD)<br>range | 9.51 (1.12) | 9.10 (1.32) | 0.24 |
|  | 7.15-10.06 | 6.47-10.86 |  |
| Gender | 24M/2F | 19M/7F |  |
| FIQ | 76-142 | 80-136 | 0.1 |
| ADOS total score | 10.89 | - |  |
| mean FD | 0.08 | 0.05 | 0.09 |
| Adolescent |  |  |  |
| Age: mean (SD)<br>range | 13.71 (1.79) | 14.01 (1.74) | 0.36 |
|  | 11.01-17.88 | 11.32-16.93 |  |
| Gender | 23M/5F | 23M/5F |  |
| FIQ | 78-132 | 80-121 | 0.52 |
| ADOS total score | 11.45 | - |  |
| mean FD | 0.07 | 0.06 | 0.3 |
| Adult |  |  |  |
| Age: mean (SD)<br>range | 24.13 (3.92) | 25.41 (5.87) | 0.32 |
|  | 18.58-39.1 | 18.59-31.78 |  |
| Gender | 14M/4F | 14M/4F |  |
| FIQ | 80-137 | 81-139 | 0.6 |
| ADOS total score | 10.82 | - |  |
| mean FD | 0.05 | 0.05 | 0.2 |

### 2.2 Dynamic FC

We used the DynamicBC toolbox (Liao et al., 2014) for Dynamic FC creation. We used a tapered sliding window length of 30s in accordance with previous studies (Allen et al., 2014; Betzel et al., 2016) and was moved in steps of 1. Consequently, a set of sliding window correlation maps was created for each participant. A Fisher’s r-to-z transformation was then applied to all the correlation maps to improve the normality of the correlation distribution. Finally, the dynamic FC variability matrix, dFCvar was calculated for each subject where D(i,j) is the standard deviation of the connection strength between ROIs i and j across the temporal windows. This matrix has also been referred to as the connection flexibility matrix in previous studies (Bassette et al., 2011, Betzel et al., 2016). Therefore, a higher D(i,j) would imply a hypervariant connection between areas i and j. Then the Network Based Statistics (NBS) method (Zalesky et al., 2010) was used to find a significantly different component (p connection < 0.01, p cluster < 0.05, 10000 iterations), both hyper-or-hypovariant in the dFCvar matrix between ASD and TD in each age group. The significant variable connections are classified into short, middle and long range connections. All the connections ie; distance between the centers of every two ROIs in the AAL atlas were listed and sorted. The shortest 33% connections are short range, highest 33% connections are long range and the other connections are defined as intermediate range. We also found the relationship between dFCvar matrix and severity score of ASD. We used the permutation method of calculating correlation. More details about the methods used as well as supplementary analysis to check for robustness of our results with different window parameters are provided in the supplementary information.

### 2.3 Modularity Maximization

Each dynamic network was treated as a layer in a multi-layer network, 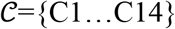. To detect the temporal evolution of modules, we maximized the multi-layer modularity, which seeks the assignment of all brain regions in all layers to modules such that the overall modularity

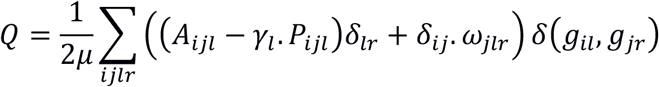

is maximized. In this expression, *Aijl* is the correlation of regions *i* and *j* in layer *l. Pijl* is the expected correlation in an appropriate null model. We used the Newman Girvan null model. The parameter, *γ*, scales the relative contribution of the expected connectivity and effectively controls the number of modules detected within a given layer. The other free parameter, *ω*, defines the weight of the inter-layer edges that link each node *i* to itself across layers. Effectively, the value of *ω* determines the consistency of multi-layer modules; when its value is large (small) relative to the intralayer links, the detected modules will tend to be more (less) similar to one another across layers. For this reason, *ω* is sometimes referred to as the *temporal resolution parameter*. For our analysis we used default values of *γ* = *ω* = 1.

Using Genlouvain Matlab toolbox (Jutla I. S. et al., 2011), we use a Louvain-like locally greedy algorithm to maximize the multi-layer modularity, *Q*. The output typically varies in each run due to near-degeneracies in the modularity landscape and stochastic elements in the optimization algorithm. For this reason, rather than focus on any single run or the consensus communities over many runs, we calculated the network flexibility, cohesion and disjointness using the iterated multilayer modularity algorithm, for 50 runs of the algorithm and then averaged those statistics over them to obtain their mean value.

### 2.4 Flexibility

The multilayer modularity maximization algorithm detected the community affiliations of each ROI at each window ie. the output is an *G = NxT* matrix where each element *(n,t)* is the community that ROI n belongs to at window t. Regional flexibility (*f*) of an ROI i, is defined as the ratio of the number of times it changed its community affiliation through the temporal windows to the possible number of community changes. It is a number between 0 and 1; where a value near 0 implies that a node’s community assignment varies very little over the course of a scan session while a value close to 1 implies that a node’s community assignment is highly variable.

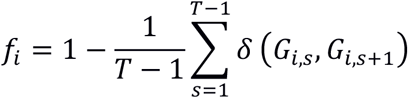

Nodes with low flexibility are considered to play a role in a temporal core while nodes with high flexibility are considered to play a role in a temporal periphery. Global flexibility (F) for a subject is calculated as the mean flexibility score across all ROIs 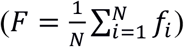.

For our analysis, we calculated both the regional and global flexibility scores separately for 50 runs of the modularity algorithm and averaged the values across runs for statistical analysis. We also correlated the regional and global flexibility scores with the subject-wise ADOS scores and the modularity of each subject.

### 2.5 Relationship between modularity and flexibility

Modularity is a measure of the excess probability of connections within the modules, relative to what is expected by chance. To calculate modularity, we first took the absolute values of each correlation and set all the diagonal elements of the correlation matrix to zero. Since fewer than 0.05% of the elements in the matrix were negative and their absolute values were relatively small, taking absolute values did not have a major effect on the results. The resulting matrix was binarized by setting the largest 10% of the edges. We used Louvain modularity algorithm (Blondel et al., 2008) to find the communities. Finally, we calculated correlation between the regional and global flexibility scores of the AAL ROIs with modularity.

### 2.6 Cohesion and Disjointness

Although flexibility characterizes the modular changes in each brain area through the scan time, it is not able to capture the method in which changes in community affiliation take place. For this, we use two network measures: Cohesion strength and node disjointness. These are complementary measures of dynamic networks. Intuitively, **node disjointedness** describes how often a node changes communities *independently* from other nodes: that is, where a node moves from community *i* to community j, and no other nodes move with it. In contrast, **node cohesion strength** describes how often a node changes communities *mutually* with other nodes. Node disjointedness is defined by the number of times a node changes communities independently, divided by the number of times a node can change communities (which is equal to the number of time windows minus unity). In contrast, node cohesion measures community changes based on the pairwise changes between nodes. It is calculated as a cohesion matrix, where the edge weight denotes the number of times a pair of nodes change to the same community together. For our study, we used a measure named cohesion strength to quantify node cohesion. It is a simple metric, analogous to dynamic degree. Cohesion strength of an ROI *i* is the sum of the row entries in the cohesion matrix. We analyzed the relation between cohesion and symptom severity by analyzing partial correlations between ADOS scores and cohesion/dispersion.

## 3. Result

### 3.1 Hypervariance of connections in ASD

We calculated the dFCvar matrix, for every subject where each connection *(i,j)* of dFCvar is the standard deviation of the connection strength between ROIs *i* and *j* across the temporal windows. We found significantly hypervariant cluster of connections in the ASD group in all age groups - children, adolescents and adults. In the children group, the cluster had 37 connections. Most of these were intra-modular DMN-DMN connections, and majority were long range connections. In the adolescents group, the cluster had 42 connections, around one third of which were intramodular and majority were short range connections. Interestingly, in adults, similar to children, we observed a majority of long range connections (**Fig 2**). Long range connections define the backbone of the functional network and often connect the hubs regions to minimize wiring and energy costs. The long range hypervariance seen in ASD children and adults could cause instability in information transmission between hubs.

**Fig 2.**
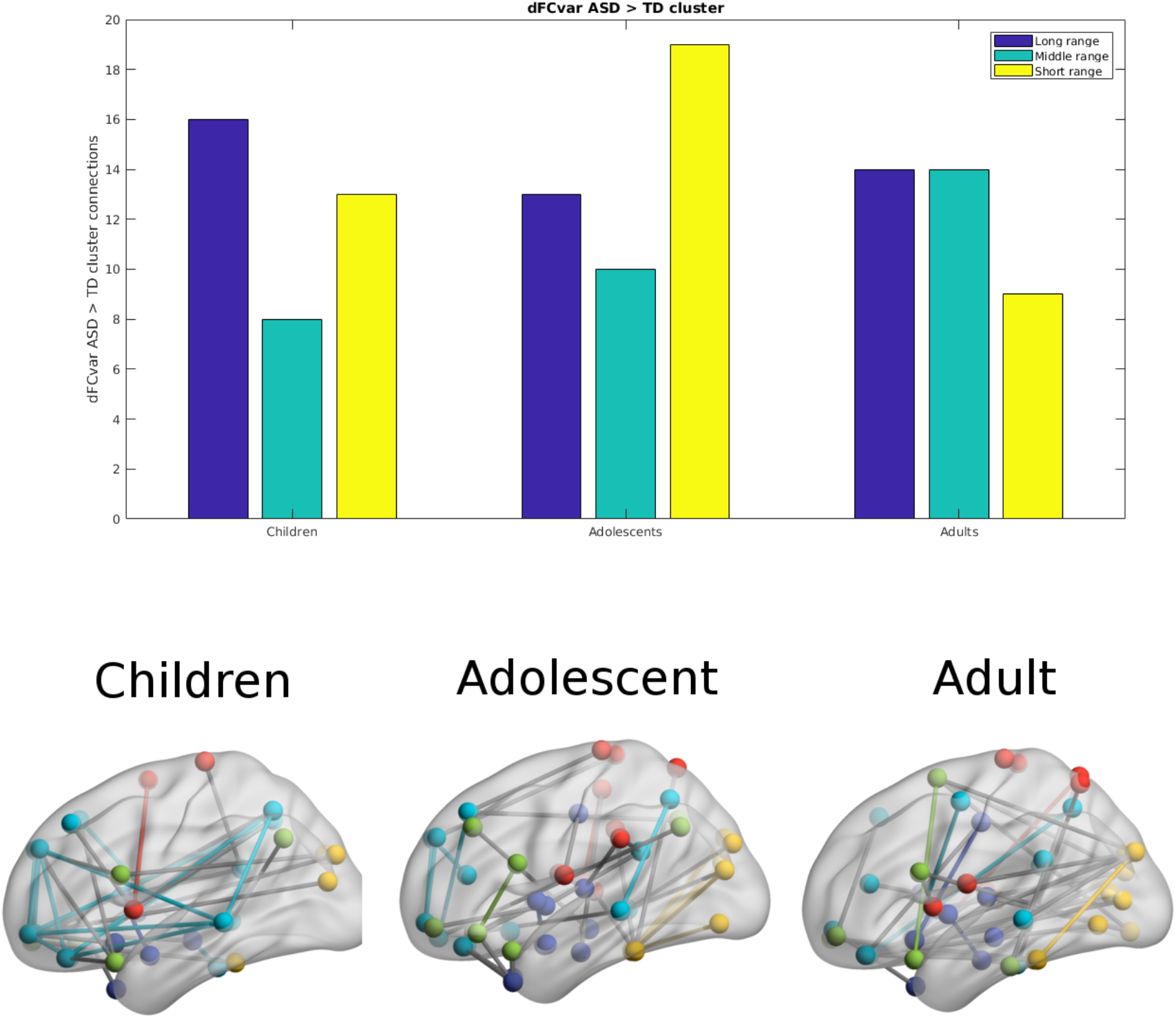
Hypervariant ASD connections in the dFCvar matrix. Majority of connections in children and adults are long range while adolescents are seen to have majority short range connections.

### 3.2 Relationship between dFCvar and ADOS scores

We found that there were 8 hypervariant connections that showed very significant correlation (r > 0.4, p <10^-5^, survived FDR correction) with ADOS score (**Table 2**). These connections did not show effect of age (one-way ANOVA, p > 0.05) indicating that this correlation was present in all age groups. These mostly included connections between inferior parietal areas and temporal regions. At the network level, we found that the mean DMN dFCvar connections showed significant correlation (r = 0.34, p < 0.05) with ADOS score as well as the mean DMN-Attention dFCvar connections (r = 0.39, p < 0.05). These too, did not show age effect with one-way ANOVA indicating that this effect is present for all age groups. However, the average functional connectivity did not show significant correlation with ADOS total scores (p values did not survive FDR correction). All within-network and between-network correlations are reported in **Supplementary materials and methods**.

**Table 1.**This shows that the inter-subject connection variability relates to symptom severity.

**Table 2.** Connections showing significant correlation (p < 10^-5^) to ADOS total score. It can be seen that in all cases, higher variability or higher dFCvar connection correlates with higher symptom severity.

| ROI_1 | ROI_2 | correlation |
| --- | --- | --- |
| Frontal_Inf_Orb_R | Parietal_Inf_L | 0.47408 |
| Parietal_Inf_L | Temporal_Mid_L | 0.42339 |
| Parietal_Inf_L | Temporal_Inf_L | 0.45008 |
| Precentral_L | Insula_R | 0.42824 |
| Precentral_L | Putamen_L | 0.48442 |
| Supp_Motor_Area_R | Insula_R | 0.44349 |
| Insula_L | Precuneus_L | 0.47195 |
| Caudate_R | Temporal_Pole_Mid_R | 0.43496 |

### 3.3 Negative correlation of flexibility with modularity

In the next analysis, we analyzed the multilayer dynamic FC using a multilayer modularity algorithm to partition the brain regions into communities across layers. For visualization, one can imagine a super adjacency matrix consisting of multiple adjacency matrices. Multilayer modularity algorithm finds communities within this super adjacency matrix. Similar to single-layer networks, brain regions assigned to the same community are more likely to be strongly connected as compared to regions assigned to different communities. Each subject had between 8 - 26 distinct communities (mean: 18.1). Based on these community assignments, we calculated the flexibility of each brain region as the fraction of times that its community assignment changed from one layer to the next (**Fig 3**).

**Fig 3.**
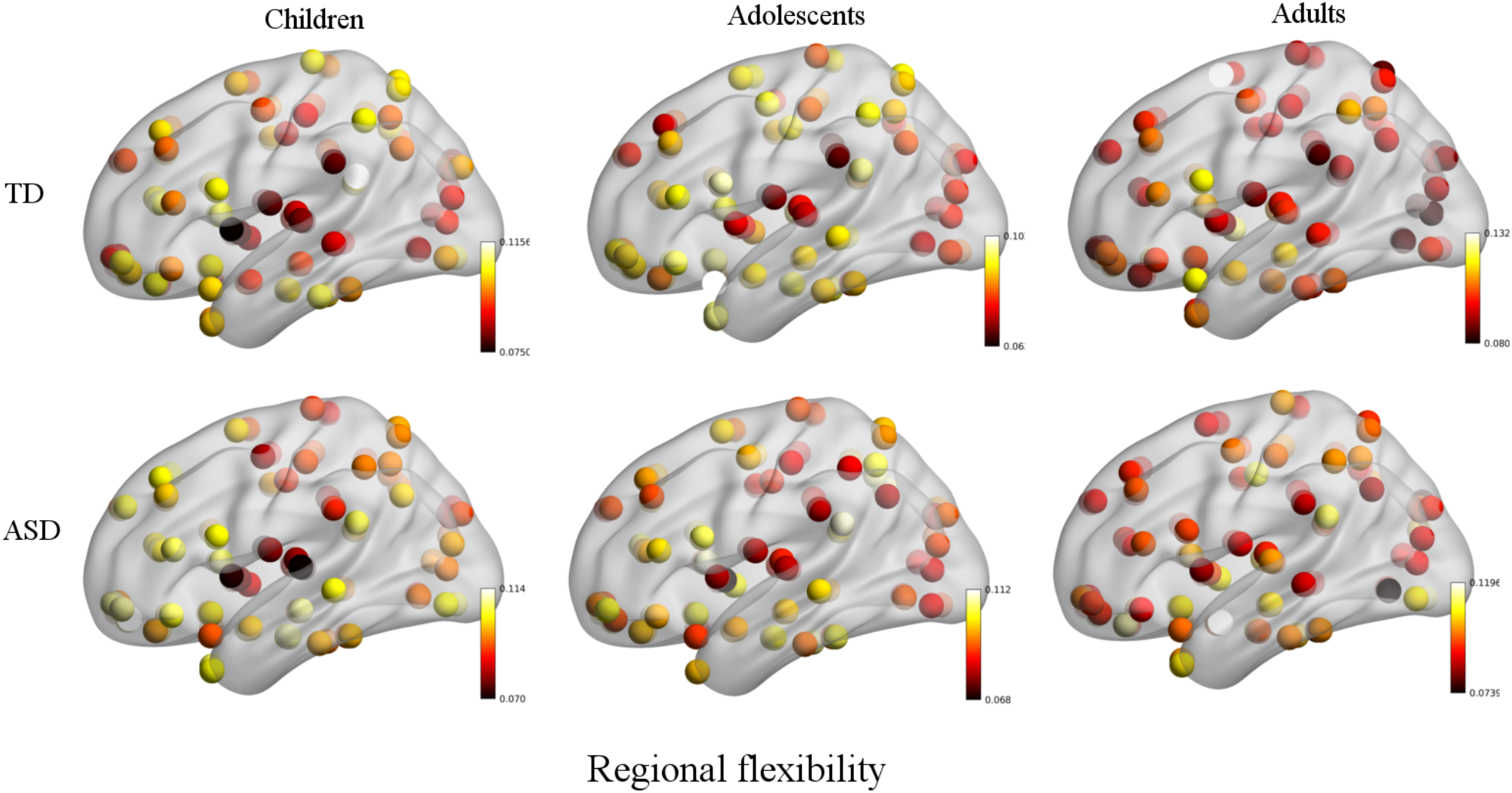
Plot of regional flexibility values for TD and ASD groups where the data is stratified into three groups: Children, adolescents and adults

Further, we calculated the modularity of static functional connectivity for each subject using Louvian algorithm and correlated it with the local and global flexibility values. In dynamic systems, different brain connectivity configurations can be interpreted as different attractor states. While modularity measures the depth of the attractor states, flexibility measures the frequency of the brain transitioning between states. Deeper or more modular states will therefore be more stable and resistant to transitions, leading to a negative correlation between modularity and flexibility (Ramos-Nuñez et al., 2017). We found that in the TD group, the mean whole-brain flexibility shows a significant (p = 0.001) correlation (r = −0.35) with the modularity (**Fig 4A**). However, ASD did not show a correlation between flexibility and modularity. On stratifying the data based on age group and then calculating the correlation, we found that in the children's age group, TD did not show any significant correlation of whole brain mean flexibility with modularity score (TD: r = −0.06, p > 0.1, ASD: r = −0.08, p > 0.1). In adolescents and adults, we found that TD showed a significant correlation (r > −0.39, p < 0.01) adult group.

**Fig 4.**
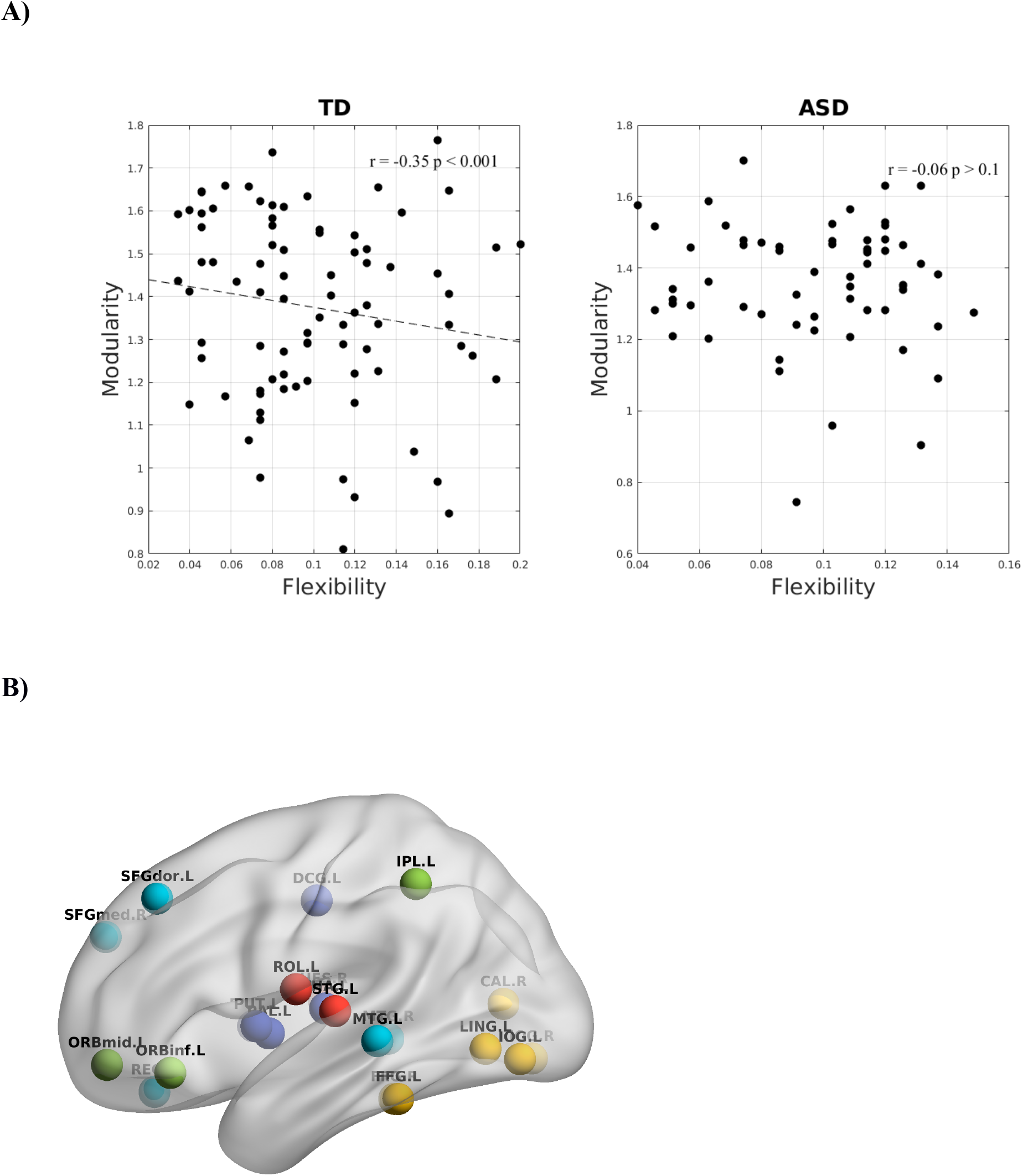
**A)** Correlation between whole-brain mean flexibility score and modularity. A significant weak negative correlation is observed in TD group which is not observed in ASD group. **B)** Areas showing significant negative correlation between local flexibility score and modularity in TD adults. None of these are significant in the ASD group.

Further, we found that there were several areas that showed a significant correlation between local flexibility score and modularity in TD while only 2 of these areas (rectus gyrus and paracentral lobule) showed this correlation in the ASD group (**Fig 4B**). The areas showing significant correlation in TD group include DMN areas: middle temporal gyrus (MTG), REC, superior frontal medial and dorsal gyrus (SFGdor, SFGmed); attention areas: inferior and middle frontal orbital areas (ORBinf, ORBmid) and inferior parietal areas (IPL); subcortical areas: putamen (PUT), PAL, thalamus (THA); visual areas: CAL, occipital inferior gyrus (IOG) and fusiform gyrus (FFG); sensorimotor/auditory areas: ROL, HES and STG, SMA.

### 3.5 Significant network level differences in dynamic network measures

To examine if the observed average changes in whole brain flexibility are driven by a biologically revelent organization of brain regions or are instead driven by randomness/noise in the whole brain, we tested the average flexibility of functional networks (DMN, Attention, Subcortical, Sensorimotor and Visual networks). We conducted a repeated measures ANOVA with functional network as a within-subject factor and with age and disease as between subject factors. We found a significant main effect of functional network (*F(38, 142)* = 3.8, *p-value* < 0.0001), while there was no significant effect of interaction of network and disease (*F(38, 142)* = 0.84, *p-value* = 0.13), or interaction of network, disease and age (*F(76, 142)* = 1.61, *p-value* = 0.16). The significant differences are listed in **Table 3**. Overall, the results primarily indicate that the visual and sensorimotor areas show least flexibility and the highest standard deviation indicating that they form the rigid temporal core. This is in contrast to DMN, Subcortical and Attention areas which have higher flexibility and lower standard deviation that form the flexible temporal periphery. **(Supplementary Tables 2, 3)**. Although there is no disease effect observed at the network level, it is possible that variation in ASD is not captured when it is considered as one group in the ANOVA analysis. Interestingly, we found that at the network level, mean visual and auditory flexibility scores show a positive correlation with the ADOS scores. However, static modularity scores did not correlate with the ADOS scores **(Fig 5A, B)**.

**Table 3.** Areas showing significant effect (p < 0.05) of age and disease on flexibility.

| ROI | Name | Abbreviation | Network | p-value | Flexibility |  |  |
| --- | --- | --- | --- | --- | --- | --- | --- |
| Disease |  |  |  |  | ASD | TD |  |
| 83 | Temporal_Pole_Sup_L | TPOsup.L | Attention | 0.036 | 0.092 | 0.107 |  |
| Age |  |  |  |  | Children | Adolescents | Adults |
| 6 | Frontal_Sup_Orb_R | ORBsup.R | DMN | 0.021 | 0.100 | 0.095 | 0.118 |
| 41 | Amygdala_L | AMYG.L | Subcortical | 0.011 | 0.095 | 0.094 | 0.116 |
| 48 | Lingual_R | LING.R | Visual | 0.043 | 0.090 | 0.074 | 0.084 |
| 53 | Occipital_Inf_L | IOG.L | Visual | 0.008 | 0.104 | 0.084 | 0.103 |
| 54 | Occipital_Inf_R | IOG.R | Visual | 0.013 | 0.108 | 0.087 | 0.105 |
| 61 | Parietal_Inf_L | IPL.L | Attention | 0.039 | 0.102 | 0.091 | 0.108 |
| 74 | Putamen_R | PUT.R | Subcortical | 0.021 | 0.089 | 0.082 | 0.105 |
| 75 | Pallidum_L | PAL.L | Subcortical | 0.001 | 0.089 | 0.089 | 0.119 |
| 77 | Thalamus_L | THA.L | Subcortical | 0.030 | 0.085 | 0.091 | 0.110 |
| 81 | Temporal_Sup_L | STG.L | Sensorimotor | 0.011 | 0.078 | 0.075 | 0.102 |

**Fig 5.**
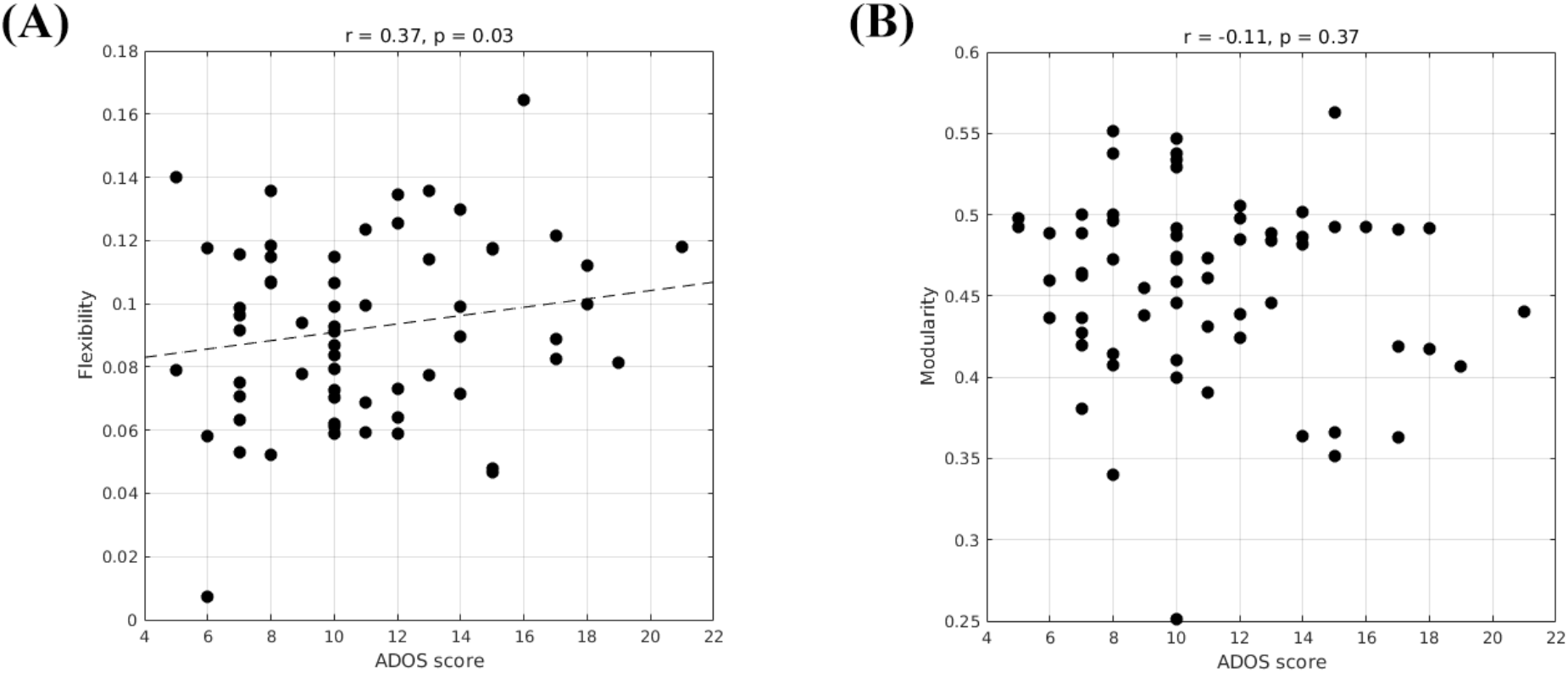
**A)** Correlation between ADOS total score and modularity **B)** Correlation between ADOS score and visual/auditory system flexibility (p < 0.05).

To further explore the differences in the dynamics at the network level, we also analysed other dynamic measures: cohesion strength and disjointness. We observed that cohesion strength shows a significant network level effect (*F* = 3.9, *p-value* < 0.01) with sensorimotor network showing high cohesion strength. There is a significant effect of network (*F* = 43.05, *p-value* < 0.0001) on node disjointness with DMN network having highest disjointness and sensorimotor, visual networks having lowest disjointness. This analysis provides insights into the organization of the dynamic resting state functional brain.

### 3.6 Effect of age and disease on dynamic metrics

We used 2-factor ANOVA with factors as disease (2 levels : TD, ASD) and age (3 levels: children, adolescents and adults) to analyze their effects on regional flexibility, cohesion and disjointness. We find that the superior temporal gyrus (TPOsup) shows significantly (F= 4.66, p = 0.03, partial eta-squared= 0.06 - medium effect size) reduced flexibility in ASD. To confirm that this difference was not driven by the difference in overall variance of the node but purely by the dynamic reconfiguration caused by the reduced flexibility, we also controlled for the mean dFCvar connections of this particular node. We still found a significant difference (p < 0.01) between ASD and TD groups. We also found several regions that show effect of age including: superior frontal orbital (ORBsup, F = 3.92, p = 0.02), PAL (F = 6.93, p = 0.001), amygdala (AMYG), cuneous (CUN), inferior occipital gyrus (IOG), left inferior parietal (IPL), angular gyrus (ANG), CAU, putamen (PUT), thalamus (THAL), SFGdor and left superior temporal (STG). Comparisons on group statistics of pallidus gyrus (periphery region) showed a significant increase in flexibility in adults as compared to both adolescents (p = 0.01) and children (p = 0.0002) while the superior frontal orbital (periphery region) shows a significant (p = 0.005) increase of flexibility in adults as compared to adolescents (**Table 3**, **Fig 6A**).

**Fig 6.**
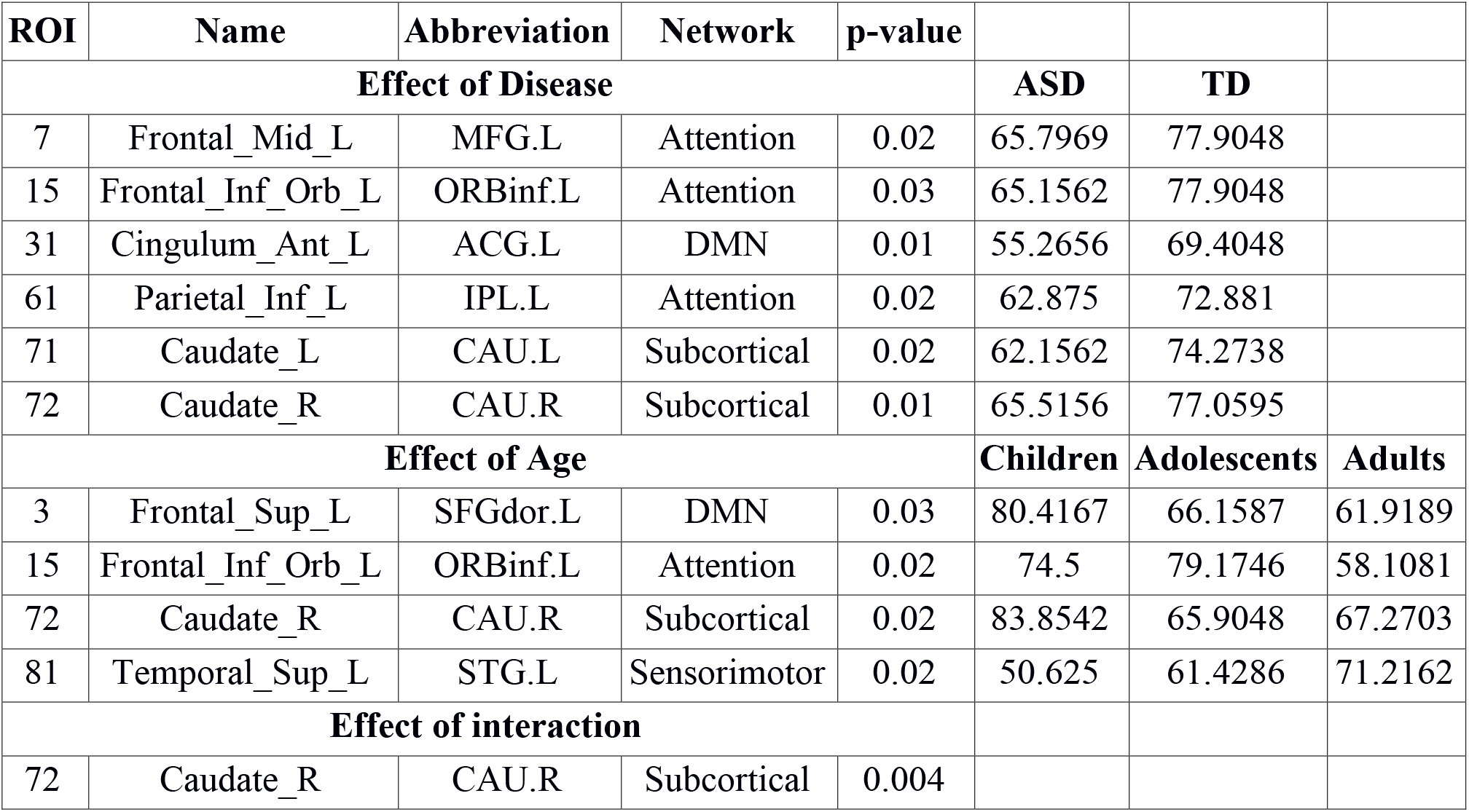
Brain plot of areas showing significant effect of age and disease on A) Flexibility, B) Cohesion strength and C) disjointness.

On analysing the effects of aging and disease on cohesion strength as well as disjointness for each individual node, we find certain nodes that show significant effect (p < 0.05). Specifically, we find that attention areas like middle frontal, inferior frontal orbital and the pareital node as well as the caudate nucleus show significant decrease in cohesion in ASD (p < 0.05, partial eta-squared ~ 0.05). While STG showed an increase in cohesion with age, areas like caudate nucleus, superior and inferior frontal areas show a general trend of decrease with age (**Fig 6B, Table 4**).

**Table 4.** Nodes showing significant effect of age, disease and interaction on cohesion strength.

| ROI | Name | Abbreviation | Network | p-value |  |  |  |
| --- | --- | --- | --- | --- | --- | --- | --- |
| Effect of Disease |  |  |  |  | ASD | TD |  |
| 7 | Frontal_Mid_L | MFG.L | Attention | 0.02 | 65.7969 | 77.9048 |  |
| 15 | Frontal_Inf_Orb_L | ORBinf.L | Attention | 0.03 | 65.1562 | 77.9048 |  |
| 31 | Cingulum_Ant_L | ACG.L | DMN | 0.01 | 55.2656 | 69.4048 |  |
| 61 | Parietal_Inf_L | IPL.L | Attention | 0.02 | 62.875 | 72.881 |  |
| 71 | Caudate_L | CAU.L | Subcortical | 0.02 | 62.1562 | 74.2738 |  |
| 72 | Caudate_R | CAU.R | Subcortical | 0.01 | 65.5156 | 77.0595 |  |
| Effect of Age |  |  |  |  | Children | Adolescents | Adults |
| 3 | Frontal_Sup_L | SFGdor.L | DMN | 0.03 | 80.4167 | 66.1587 | 61.9189 |
| 15 | Frontal_Inf_Orb_L | ORBinf.L | Attention | 0.02 | 74.5 | 79.1746 | 58.1081 |
| 72 | Caudate_R | CAU.R | Subcortical | 0.02 | 83.8542 | 65.9048 | 67.2703 |
| 81 | Temporal_Sup_L | STG.L | Sensorimotor | 0.02 | 50.625 | 61.4286 | 71.2162 |
| Effect of interaction |  |  |  |  |  |  |  |
| 72 | Caudate_R | CAU.R | Subcortical | 0.004 |  |  |  |

The rolandic operculum gyrus, lingual and occipital gyrus as well as the medial frontal orbital show a significant increase in disjointness in ASD (p < 0.05, partial eta-squared ~ 0.06). Further, several nodes in DMN and Sensorimotor network show significant decrease in node doisjointness with aging (p < 0.05, partial eta-squared ~ 0.07) (**Fig 6C, Table 5**).

**Table 5.** Nodes showing significant effect of age, disease and interaction on node disjointness. Overall, ASD shows an increase in disjointness and a decrease in node cohesion strength.

| ROI | Name | Abbreviation | Network | p-value | Node disjointness |  |  |
| --- | --- | --- | --- | --- | --- | --- | --- |
| Effect of Disease |  |  |  |  | ASD | TD |  |
| 17 | Rolandic_Oper_L | ROL.L | Sensorimotor | 0.01 | 0.0039 | 0.00217 |  |
| 26 | Frontal_Med_Orb_R | ORBsupmed.R | DMN | 0.04 | 0.00482 | 0.00340 |  |
| 47 | Lingual_L | LING.L | Visual | 0.04 | 0.00473 | 0.00299 |  |
| 53 | Occipital_Inf_L | IOG.L | Visual | 0.01 | 0.00741 | 0.00455 |  |
| 58 | Postcentral_R | PoCG.R | Sensorimotor | 0.009 | 0.00669 | 0.00414 |  |
| Effect of Age |  |  |  |  | Children | Adolescents | Adults |
| 3 | Frontal_Sup_L | SFGdor.L | DMN | 0.01 | 0.0075 | 0.004 | 0.0055 |
| 4 | Frontal_Sup_R | SFGdor.R | DMN | 0.04 | 0.0051 | 0.0061 | 0.003 |
| 18 | Rolandic_Oper_R | ROL.R | Sensorimotor | 0.001 | 0.0019 | 0.002 | 0.0052 |
| 32 | Cingulum_Ant_R | ACG.R | DMN | 0.03 | 0.0067 | 0.0063 | 0.0033 |
| 46 | Cuneus_R | CUN.R | Visual | 0.005 | 0.00321 | 0.0029 | 0.0060 |
| 60 | Parietal_Sup_R | SPG.R | Sensorimotor | 0.03 | 0.0063 | 0.0056 | 0.0027 |
| 75 | Pallidum_L | PAL.L | Subcortical | 0.03 | 0.0036 | 0.0054 | 0.0074 |
| 76 | Pallidum_R | PAL.R | Subcortical | 0.03 | 0.0066 | 0.009 | 0.006 |

### 3.7 Correlation of dynamic metrics (local dynamics) with ADOS scores

To analyze the relation between symptom severity and dynamic measures – flexibility, cohesion strength and disjointness, we found their correlations with the ADOS scores.While we did not find a significant correlation between mean whole brain flexibility scores with ADOS scores, at the functional network level, we found the mean flexibility score by averaging the individual scores of each node in a functional network. We found that an increase in flexibility score of visual system areas positively correlated with the ADOS Total symptom severity score (r = 0.37, p < 0.01); ADOS Social score (r = 0.38, p < 0.01). We did not find any significant correlation with the other functional networks. At the level of individual ROIs, we found that the flexibility of several visual and auditory areas showed positive correlation with ADOS total and communication scores. These results are summarized in **Supplementary Table 4** and the these areas are plotted in **Supplementary Fig 1.** Further, on dividing the subjects into 3 age groups - children, adolescents and adults, we found that adolescents and adults show a very strong positive correlation of ADOS social score and flexibility in rigid regions. Children show a strong negative correlation of ADOS score and flexibility of DMN region - which is a flexible periphery region (**Table 6**).

**Table 6.** Areas showing significant correlation between region flexibility and ADOS social scores in each age group.

| Node flexibility |  |  |  |  |  |
| --- | --- | --- | --- | --- | --- |
| ROI | Abbreviation | Network | Name | r | p |
| Children |  |  |  |  |  |
| 50 | SOG.R | Visual | Occipital_Sup_R | 0.50199 | 0.01056 |
| 86 | MTG.R | DMN | Temporal_Mid_R | -0.41371 | 0.039801 |
| Adolescents |  |  |  |  |  |
| 17 | ROL.L | Sensorimotor | Rolandic_Oper_L | 0.45692 | 0.024791 |
| 43 | CAL.L | Visual | Calcarine_L | 0.52458 | 0.0084972 |
| 45 | CUN.L | Visual | Cuneus_L | 0.41229 | 0.04528 |
| 46 | CUN.R | Visual | Cuneus_R | 0.47192 | 0.019898 |
| 48 | LING.R | Visual | Lingual_R | 0.53131 | 0.0075468 |
| 50 | SOG.R | Visual | Occipital_Sup_R | 0.49459 | 0.014015 |
| 72 | CAU.R | Subcortical | Caudate_R | 0.46562 | 0.021849 |
| Adults |  |  |  |  |  |
| 21 | OLF.L | Sensorimotor | Olfactory_L | 0.69289 | 0.0041875 |
| 43 | CAL.L | Visual | Calcarine_L | 0.61222 | 0.015269 |
| 45 | CUN.L | Visual | Cuneus_L | 0.55795 | 0.030667 |

On analyzing correlation between the dynamic properties of node disjointness as well as node cohesion strength with ADOS scores, we find that for adults, the mean node disjointness i.e. the average over the whole brain; shows a significant positive correlation (r = 0.45, p < 0.01) with the symptom severity ADOS scores. We also find several nodes that show significant correlation of cohesion strength with the ADOS scores. We have listed the details of the correlations with ADOS scores in **Tables 7, 8**.

**Table 7.**
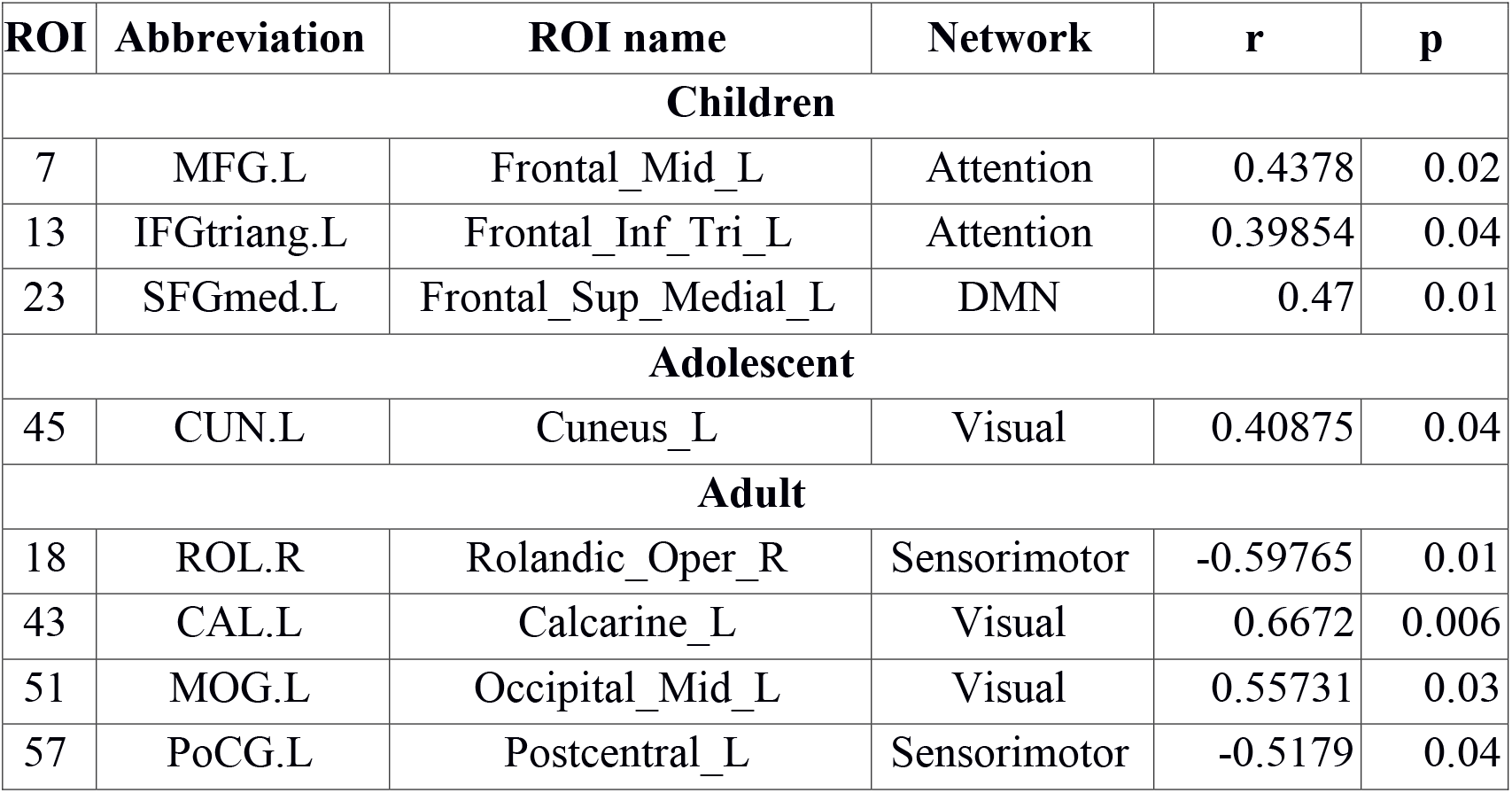
Nodes showing significant correlation of node cohesion strength with ADOS social score.

| ROI | Abbreviation | ROI name | Network | r | p |
| --- | --- | --- | --- | --- | --- |
| Children |  |  |  |  |  |
| 7 | MFG.L | Frontal_Mid_L | Attention | 0.4378 | 0.02 |
| 13 | IFGtriang.L | Frontal_Inf_Tri_L | Attention | 0.39854 | 0.04 |
| 23 | SFGmed.L | Frontal_Sup_Medial_L | DMN | 0.47 | 0.01 |
| Adolescent |  |  |  |  |  |
| 45 | CUN.L | Cuneus_L | Visual | 0.40875 | 0.04 |
| Adult |  |  |  |  |  |
| 18 | ROL.R | Rolandic_Oper_R | Sensorimotor | -0.59765 | 0.01 |
| 43 | CAL.L | Calcarine_L | Visual | 0.6672 | 0.006 |
| 51 | MOG.L | Occipital_Mid_L | Visual | 0.55731 | 0.03 |
| 57 | PoCG.L | Postcentral_L | Sensorimotor | -0.5179 | 0.04 |

**Table 8.** Nodes showing significant correlation of node disjointness with ADOS social score.

| ROI | Abbreviation | ROI name | Network | r | p |
| --- | --- | --- | --- | --- | --- |
| **Children** |  |  |  |  |  |
| 14 | IFGtriang.R | Frontal_Inf_Tri_R | Attention | 0.45644 | 0.02 |
| 34 | DCG.R | Cingulum_Mid_R | Subcortical | 0.56819 | 0.003 |
| 84 | TPOsup.R | Temporal_Pole_Sup_R | Sensorimotor | 0.5094 | 0.009 |
| **Adolescent** |  |  |  |  |  |
| 24 | SFGmed.R | Frontal_Sup_Medial_R | DMN | -0.57119 | 0.003 |
| 46 | CUN.R | Cuneus_R | Visual | 0.52479 | 0.008 |
| 48 | LING.R | Lingual_R | Visual | 0.54853 | 0.005 |
| 82 | STG.R | Temporal_Sup_R | Sensorimotor | 0.47184 | 0.01 |
| 86 | MTG.R | Temporal_Mid_R | DMN | -0.55651 | 0.004 |
| **Adult** |  |  |  |  |  |
| 42 | AMYG.R | Amygdala_R | Subcortical | 0.52244 | 0.04 |
| 50 | SOG.R | Occipital_Sup_R | Visual | 0.57317 | 0.02 |

Overall, we see that while increase in cohesion strength in significant sensorimotor regions correlates negatively with ADOS scores for ASD adults, the affected visual, attention and DMN areas correlate positively. DMN areas show negative correlation of disjointness with ADOS scores and vice versa for sensorimotor areas.

## 4. Discussion

Autism is a neurodevelopmental disorder characterized by altered neural network dynamics. The presence of alterations in the time-varying configuration of the functional connectome and their implications on symptom severity are yet to be studied. To address this gap in knowledge, our study applied innovative methods from dynamic network neuroscience to resting-state fMRI data.

In our first analysis, we found that mean dFCvar (connection flexibility) between the Attention and DMN networks is positively correlated with the ADOS scores. This indicates that the inter-subject variability is related to the symptom severity. A previous study showed that although during a task, higher dynamic connection variability between Attention and DMN is associated with better cognitive performance, the opposite is true for resting-state Attention–DMN dFCvar (Douw et al., 2016). Similarly, Lin et al. (2016) reported that higher variability in the connection strength of PCC to other DMN areas in the resting-state is related to slower reaction times on a subsequent attention task. The hypervariance in ASD could lead to the globally disconnected state in its dynamics that has been reported in a previous study (Rashid et al., 2018). These results taken together indicate that there could be a relation between the atypical hypervariance in ASD which leads to increase in ADOS score and decrease in cognitive performance.

In the next analysis, we explored dynamic network measures – like network flexibility, cohesion and disjointness to understand changes in functional connectivity that occur over time. Recent work in the field of network flexibility has centered around the relationship between task-performance or task-sentiment analysis with dynamic reconfiguration of the brain (Park et al., 2017; Telesford et al., 2017) especially in the memory areas (Douw et al., 2015). Intra-subject changes in flexibility have been associated with mood (Betzel et al., 2017), while inter-subject differences have been linked to learning (Bassett et al., 2011), working memory performance (Braun et al., 2015), and reinforcement learning (Gerraty et al., 2018). The metric has also been found to correlate with schizophrenia risk, and is altered by an NMDA-receptor antagonist (Braun et al., 2016). Flexibility also has age-based variation (Schlesinger et al., 2017). Recent work has established that there is a significant negative correlation of flexibility and modularity using task-based fMRI studies (Ramos-Nuñez et al., 2017). In our study, we found that such a correlation exists even in the resting-state functional brain as well. We found that while dynamic flexibility correlates negatively with the static metric of modularity in TD, such a correlation does not exist in ASD (r = −0.1, p > 0.1) . However, both TD and ASD showed a range of flexibilities, indicating that the whole-brain flexibility remains intact. This indicates that the modular re-organization with variations in flexibility is not observed in ASD. It is possible that there are network/nodal changes that compensate for the change in flexibility. We also found that modularity scores did not correlate with the symptom severity ADOS score while network-based flexibility scores showed a significant positive/negative correlation. This further indicates that static network measures are unable to capture the underlying variability in ASD with respect to the ADOS scores. In contrast, dynamic network measures have significant predictive power of ADOS scores.

We found that in the resting-state functional brain, regions with lower flexibility are those involved in visual, hearing and motor processes while those with higher flexibility are those typically associated with the default mode network, cognitive control and executive function. This organization has been previously reported in task-based fMRI studies (Cole et al., 2013; Braun et al., 2015; Mattar et al., 2015; Schlesinger et al., 2017). Cole et al (2013) showed that a flexible frontoparietal architecture across tasks was associated with execution of multiple cognitive tasks. de Lacy et al (2017) reported that the number of state transitions were reduced in ASD as compared to TD, due to disruptions in the fronto-pareital network and impaired state transitions in the cingulo-opercular systems. In our study, we found that there is reduced flexibility of periphery regions (DMN, subcortical and attention areas) in ASD which could impair the state transitions.

Although the network flexibility did not show disease or age effect, we found regions (nodes) that show their independent effect. Significant effect of disease is observed in TPOsup, which is an important area linked to verbal and nonverbal communication identified to show abnormal behaviour in autism. It is a key periphery region which shows increased flexibility in TD group. The decrease of flexibility of TPOsup could contribute to the nature of atypical verbal behaviour observed in ASD. On analyzing effect of age, disease and their interaction on node cohesion and disjointness, we observed that ASD showed an atypical decrease in node cohesion strength and an increase in disjointness. Telesford and colleagues (Telesford et al., 2017) found that node cohesion strength showed positive correlation with performance in a task. Our results suggest that for resting-state dynamics, sensorimotor and visual regions show high cohesion strength while DMN shows relatively high disjointness. Further, while sensorimotor cohesion strength correlates negatively (r = −0.6) with ADOS symptom severity score, for significantly affected nodes in other networks, the correlation is positive (r ~ 0.5). The findings suggest that while sensorimotor areas are least flexible, they are required to be less cohesive as well.

## 5. Conclusion

To the best of our knowledge, this is one of the first studies to use dynamic modularity metrics like flexibility and cohesion to study altered dynamics in ASD and TD. Our study differs in several aspects from previously reported dynamic whole-brain network level investigations in ASD (Chen et al., 2017; de Lacy et al., 2017; Rashid et al., 2018). Most of the previous studies on dynamic functional connectivity in ASD combined sliding window analysis with k-means clustering to identify common brain states among all subjects (both ASD and TD). On the other hand, Watanabe and Rees et al (2017) concatenated the timeseries of ASD subjects to characterize the energy landscape of ASD. However, in our study, we have captured the variability among subjects by performing dynamic analysis on each individual subject. We uncovered the effect of development, disease and their interactions on the dynamic metrics.

We found very high correlations of ADOS score with the flexibility and disjointness of Sensorimotor regions. This is indicative of the importance of maintaining the cohesion and rigidity of motor cortex regions in TD. It can be noted that in our study, static modularity does not show a correlation with ADOS scores. Using temporal modularity metrics for dynamic analysis is a recent approach and lacks standardization across clinical studies. Whole brain connectivity anomaly and differences are highly sensitive to the parameters used while partitioning the BOLD timeseries into instantaneous dynamic FC matrices. To study the robustness of our results, we also repeated the dFCvar analysis using different window parameters and found consistent results (Supplementary analysis). Further, it is also extremely important to correct for head motion as it is an important confounding factor. Several studies have shown that not correcting for it sufficiently can lead to spurious instantaneous connections, either increased or decreased. In our study, we have exercised scrutiny by using samples from a single site which are corrected for motion artifacts and considered samples with low framewise displacement that were approved by manual functional QA raters.

Overall, this study provides insights into the patterns observed in the functional brain systems of TD and ASD from a dynamic perspective. This can further be extended using a longitudinal design, including a larger subject pool and combining structural data as well.

## 6. Acknowledgements

DR is supported by the Department of Biotechnology DBT Ramalingaswami fellowship (BT/RLF/Re-entry/07/2014) and DST Cognitive Science Research Initiative grant (SR/CSRI/21/2016). PKV acknowledges financial support from the Early Career Research Award Scheme (ECR/2016/000488), Science and Engineering Research Board, DST, India.

## 7. Disclosure Statement

No competing financial interests exist.

## Supplementary Information

### Correlation calculation

1. Correlation step: for each connection in the dynamic FC variability matrix, Pearson correlation coefficients were calculated between the dFCvar and ADOS total score. The number of significantly positively correlated connections (P < 0.05) was summed.
2. Permutation step: the subjects were shuffled to create a random dataset, then the Pearson correlation coefficients between the dFCvar of each connection and ADOS total score were calculated in the shuffled dataset. The number of significantly positively correlated connections (P < 0.05) was summed. This step was repeated 5000 times and 5000 significantly positively correlated connections at random were obtained. p value of the hypervariant cluster was calculated by the times in which the random correlated connections were more than the actual correlated connections divided by 5000.

### Robustness with window parameters

To analyze the robustness of our analysis with window parameters, we repeated the dynamic FC analysis with a window size of 40s and step size of 2. We found that for children, adolescents and adults, ASD showed significantly greater variability (p < 0.05) as compared to TD. For children, the cluster size is 44 connections, with majority connections (19 connections) being long range. For adolescents, the cluster size is 34 connections, with majority as short range (15 connections). For adults, the cluster size is 52 connections with 19 long range and 20 middle range connections. On correlating dFCvar with ADOS scores, we found that the DMN-DMN as well as DMN-Attention mean dFCvar showed significant positive correlation (r > 0.3, p < 0.05). Further, 8 connections were identified that showed high correlation with ADOS score (r > 0.5, p < 0.05). Overall, the distribution of the hypervariant connections as well as ADOS correlation is consistent with our reported results.

### Supplementary tables

**Supplementary Table 1.** Between networks and within network correlation of dFCvar with ADOS scores.

| Network_1 | Network_2 | correlation | p |
| --- | --- | --- | --- |
| DMN | DMN | 0.34 | 0.04 |
|  | visual | -0.05 | 0.66 |
|  | attention | 0.39 | 0.01 |
|  | subcortical | 0.15 | 0.22 |
|  | sensorimotor | 0.16 | 0.2 |
| Visual | visual | 0.22 | 0.07 |
|  | attention | 0.06 | 0.58 |
|  | subcortical | 0.05 | 0.67 |
|  | sensorimotor | 0.17 | 0.15 |
| attention | attention | 0.2 | 0.1 |
|  | subcortical | 0.17 | 0.16 |
|  | sensorimotor | 0.19 | 0.12 |
| subcortical | subcortical | 0.24 | 0.051 |
|  | sensorimotor | 0.24 | 0.052 |
| sensorimotor | sensorimotor | 0.23 | 0.06 |

**Supplementary Table 2.**
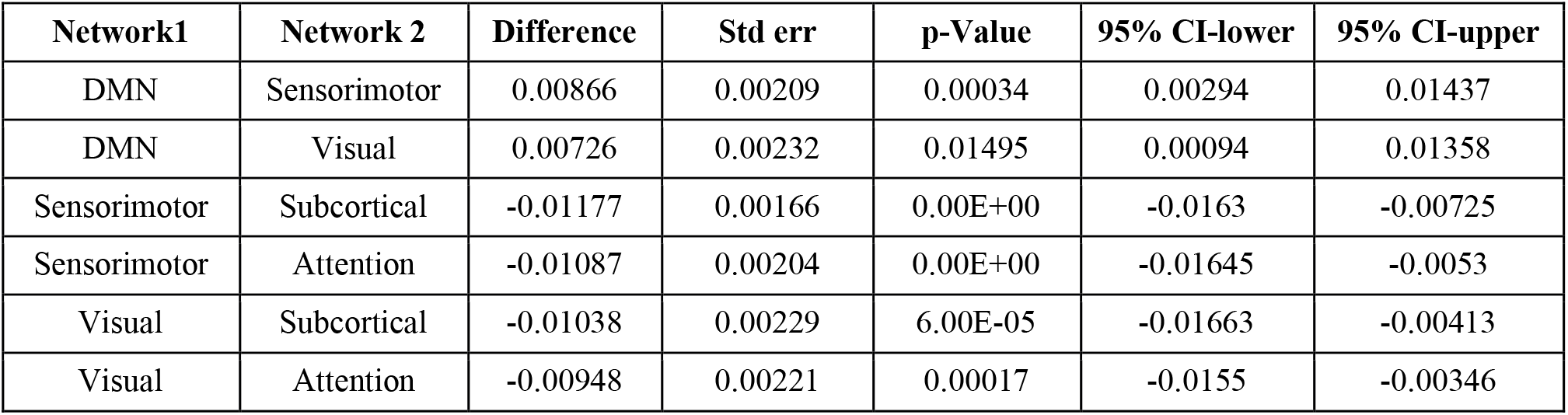
Statistics of difference in flexibility between networks for networks showing significantchanges.

**Supplementary Table 3.** Group statistics of flexibility score of each network.

| Network | Mean | Std dev |
| --- | --- | --- |
| Default Mode | 0.096 | 0.023 |
| Sensorimotor | 0.087 | 0.027 |
| Visual | 0.089 | 0.027 |
| Subcortical | 0.099 | 0.024 |
| Attention | 0.098 | 0.022 |

**Supplementary Table 4.** Areas showing significant correlation of regional flexibility score with ADOS score.

| ROI # | Name | Network | Abbreviation | r | p |
| --- | --- | --- | --- | --- | --- |
| ADOS COMMUNICATION score |  |  |  |  |  |
| 17 | Rolandic_Oper_L | Sensorimotor | ROL.L | 0.31 | 0.012 |
| 22 | Olfactory_R | Sensorimotor | OLF.R | 0.25 | 0.041 |
| 89 | Temporal_Inf_L | Attention | ITG.L | 0.29 | 0.017 |
| ADOS SOCIAL score |  |  |  |  |  |
| 22 | Olfactory_R | Sensorimotor | OLF.R | 0.32 | 0.01 |
| 26 | Frontal_Med_Orb_R | DMN | ORBsupmed.R | 0.28 | 0.02 |
| 43 | Calcarine_L | Visual | CAL.L | 0.35 | 0.0038 |
| 44 | Calcarine_R | Visual | CAL.R | 0.26 | 0.037 |
| 45 | Cuneus_L | Visual | CUN.L | 0.29 | 0.018 |
| 49 | Occipital_Sup_L | Visual | SOG.L | 0.28 | 0.021 |
| 50 | Occipital_Sup_R | Visual | SOG.R | 0.45 | 0.00014 |
| 82 | Temporal_Sup_R | Sensorimotor | STG.R | 0.29 | 0.049 |
| 89 | Temporal_Inf_L | Attention | ITG.L | 0.27 | 0.027 |
| ADOS TOTAL score |  |  |  |  |  |
| 17 | Rolandic_Oper_L | Sensorimotor | ROL.L | 0.27 | 0.026 |
| 22 | Olfactory_R | Sensorimotor | OLF.R | 0.33 | 0.0075 |
| 26 | Frontal_Med_Orb_R | DMN | ORBsupmed.R | 0.26 | 0.038 |
| 43 | Calcarine_L | Visual | CAL.L | 0.32 | 0.0096 |
| 45 | Cuneus_L | Visual | CUN.L | 0.29 | 0.018 |
| 46 | Cuneus_R | Visual | CUN.R | 0.25 | 0.045 |
| 49 | Occipital_Sup_L | Visual | SOG.L | 0.25 | 0.046 |
| 50 | Occipital_Sup_R | Visual | SOG.R | 0.42 | 0.00043 |
| 89 | Temporal_Inf_L | Attention | ITG.L | 0.31 | 0.011 |

**Supplementary Figure 1.**
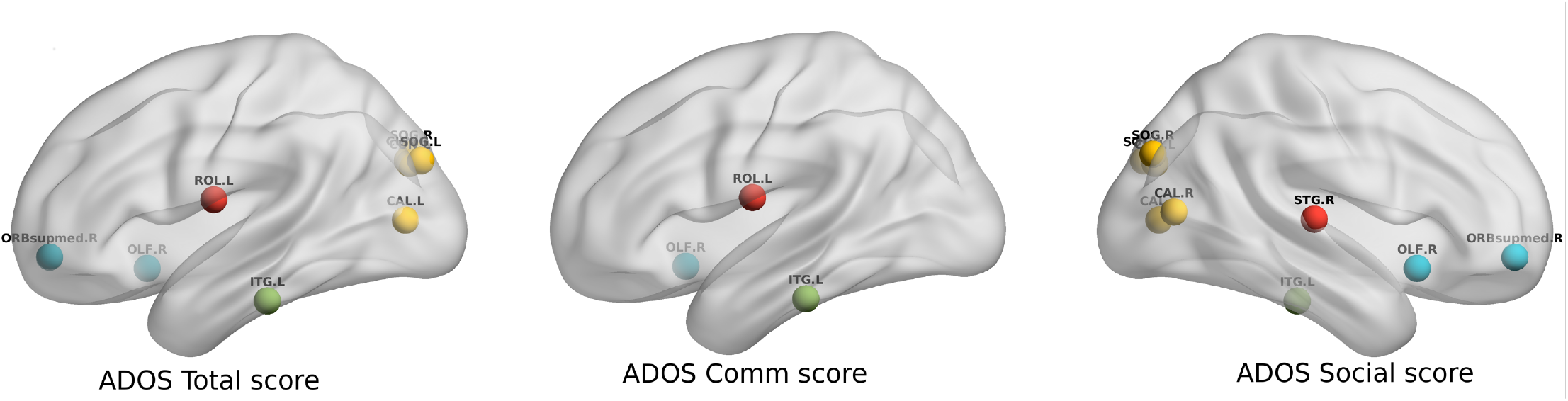
Areas showing significant (p < 0.05) correlation of flexibility score with ADOS scores.

## References

Allen, E.A., Damaraju, E., Plis, S.M., Erhardt, E.B., Eichele, T., Calhoun, V.D., 2014. Tracking Whole-Brain Connectivity Dynamics in the Resting State. Cerebral Cortex 24, 663–676. https://doi.org/10.1093/cercor/bhs352

Bassett, D.S., Wymbs, N.F., Porter, M.A., Mucha, P.J., Carlson, J.M., Grafton, S.T., 2011. Dynamic reconfiguration of human brain networks during learning. Proceedings of the National Academy of Sciences 108, 7641–7646. https://doi.org/10.1073/pnas.1018985108

Betzel, R.F., Fukushima, M., He, Y., Zuo, X.-N., Sporns, O., 2016. Dynamic fluctuations coincide with periods of high and low modularity in resting-state functional brain networks. NeuroImage 127, 287–297. https://doi.org/10.1016/j.neuroimage.2015.12.001

Betzel, R.F., Satterthwaite, T.D., Gold, J.I., Bassett, D.S., 2017. Positive affect, surprise, and fatigue are correlates of network flexibility. Scientific Reports 7. https://doi.org/10.1038/s41598-017-00425-z

Blondel, V.D., Guillaume, J.-L., Lambiotte, R., Lefebvre, E., 2008. Fast unfolding of communities in large networks. Journal of Statistical Mechanics: Theory and Experiment 2008, P10008. https://doi.org/10.1088/1742-5468/2008/10/P10008

Braun, U., Schäfer, A., Bassett, D.S., Rausch, F., Schweiger, J.I., Bilek, E., Erk, S., Romanczuk-Seiferth, N., Grimm, O., Geiger, L.S., Haddad, L., Otto, K., Mohnke, S., Heinz, A., Zink, M., Walter, H., Schwarz, E., Meyer-Lindenberg, A., Tost, H., 2016. Dynamic brain network reconfiguration as a potential schizophrenia genetic risk mechanism modulated by NMDA receptor function. Proceedings of the National Academy of Sciences 113, 12568–12573. https://doi.org/10.1073/pnas.1608819113

Braun, U., Schäfer, A., Walter, H., Erk, S., Romanczuk-Seiferth, N., Haddad, L., Schweiger, J.I., Grimm, O., Heinz, A., Tost, H., Meyer-Lindenberg, A., Bassett, D.S., 2015. Dynamic reconfiguration of frontal brain networks during executive cognition in humans. Proceedings of the National Academy of Sciences 112, 11678–11683. https://doi.org/10.1073/pnas.1422487112

Cameron, C., Yassine, B., Carlton, C., Francois, C., Alan, E., András, J., Budhachandra, K., John, L., Qingyang, L., Michael, M., Chaogan, Y., Pierre, B., 2013. The Neuro Bureau Preprocessing Initiative: open sharing of preprocessed neuroimaging data and derivatives. Frontiers in Neuroinformatics 7. https://doi.org/10.3389/conf.fninf.2013.09.00041

Chang, C., Glover, G.H., 2010. Time–frequency dynamics of resting-state brain connectivity measured with fMRI. NeuroImage 50, 81–98. https://doi.org/10.1016/j.neuroimage.2009.12.011

Chen, Guangyu, Zhang, H.-Y., Xie, C., Chen, Gang, Zhang, Z.-J., Teng, G.-J., Li, S.-J., 2013. Modular reorganization of brain resting state networks and its independent validation in Alzheimer’s disease patients. Frontiers in Human Neuroscience 7. https://doi.org/10.3389/fnhum.2013.00456

Chen, Heng, Nomi, J.S., Uddin, L.Q., Duan, X., Chen, Huafu, 2017. Intrinsic functional connectivity variance and state-specific under-connectivity in autism: State-Related Functional Connectivity in Autism. Human Brain Mapping 38, 5740–5755. https://doi.org/10.1002/hbm.23764

Cole, M.W., Reynolds, J.R., Power, J.D., Repovs, G., Anticevic, A., Braver, T.S., 2013. Multi-task connectivity reveals flexible hubs for adaptive task control. Nature Neuroscience 16, 1348–1355. https://doi.org/10.1038/nn.3470

Damaraju, E., Allen, E.A., Belger, A., Ford, J.M., McEwen, S., Mathalon, D.H., Mueller, B.A., Pearlson, G.D., Potkin, S.G., Preda, A., Turner, J.A., Vaidya, J.G., van Erp, T.G., Calhoun, V.D., 2014. Dynamic functional connectivity analysis reveals transient states of dysconnectivity in schizophrenia. NeuroImage: Clinical 5, 298–308. https://doi.org/10.10167j.nicl.2014.07.003

de Lacy, N., Doherty, D., King, B.H., Rachakonda, S., Calhoun, V.D., 2017. Disruption to control network function correlates with altered dynamic connectivity in the wider autism spectrum. NeuroImage: Clinical 15, 513–524. https://doi.org/10.1016Zj.nicl.2017.05.024

Deco, G., Ponce-Alvarez, A., Mantini, D., Romani, G.L., Hagmann, P., Corbetta, M., 2013. Resting-State Functional Connectivity Emerges from Structurally and Dynamically Shaped Slow Linear Fluctuations. Journal of Neuroscience 33, 11239–11252. https://doi.org/10.1523/JNEUROSCI.1091-13.2013

Douw, L., Leveroni, C.L., Tanaka, N., Emerton, B.C., Cole, A.C., Reinsberger, C., Stufflebeam, S.M., 2015. Loss of Resting-State Posterior Cingulate Flexibility Is Associated with Memory Disturbance in Left Temporal Lobe Epilepsy. PLOS ONE 10, e0131209. https://doi.org/10.1371/journal.pone.0131209

Douw, L., Wakeman, D.G., Tanaka, N., Liu, H., Stufflebeam, S.M., 2016. State-dependent variability of dynamic functional connectivity between frontoparietal and default networks relates to cognitive flexibility. Neuroscience 339, 12–21. https://doi.org/10.1016/j.neuroscience.2016.09.034

Falahpour, M., Thompson, W.K., Abbott, A.E., Jahedi, A., Mulvey, M.E., Datko, M., Liu, T.T., Müller, R.-A., 2016. Underconnected, But Not Broken? Dynamic Functional Connectivity MRI Shows Underconnectivity in Autism Is Linked to Increased Intra-Individual Variability Across Time. Brain Connectivity 6, 403–414. https://doi.org/10.1089/brain.2015.0389

Garcia, J.O., Ashourvan, A., Muldoon, S., Vettel, J.M., Bassett, D.S., 2018. Applications of Community Detection Techniques to Brain Graphs: Algorithmic Considerations and Implications for Neural Function. Proceedings of the IEEE 106, 846–867. https://doi.org/10.1109/JPROC.2017.2786710

Gerraty, R.T., Davidow, J.Y., Foerde, K., Galvan, A., Bassett, D.S., Shohamy, D., 2018. Dynamic flexibility in striatal-cortical circuits supports reinforcement learning. The Journal of Neuroscience 2084–17. https://doi.org/10.1523/JNEUROSCI.2084-17.2018

Hahamy, A., Behrmann, M., Malach, R., 2015. The idiosyncratic brain: distortion of spontaneous connectivity patterns in autism spectrum disorder. Nature Neuroscience 18, 302–309. https://doi.org/10.1038/nn.3919

Harlalka, V., Raju, B.S., Vinod, P.K., Roy, D., 2018. Age, disease and their interaction effects on intrinsic connectivity of children and adolescents in Autism Spectrum Disorder using functional connectomics. Brain Connectivity. https://doi.org/10.1089/brain.2018.0616

Jutla I. S., Jeub L. G. S, Mucha P. J., 2011. A generalized Louvain method for community detection implemented in matlab.

Lee, M.H., Smyser, C.D., Shimony, J.S., 2013. Resting-State fMRI: A Review of Methods and Clinical Applications. American Journal of Neuroradiology 34, 1866–1872. https://doi.org/10.3174/ajnr.A3263

Liao, W., Wu, G.-R., Xu, Q., Ji, G.-J., Zhang, Z., Zang, Y.-F., Lu, G., 2014. DynamicBC : A MATLAB Toolbox for Dynamic Brain Connectome Analysis. Brain Connectivity 4, 780–790. https://doi.org/10.1089/brain.2014.0253

Lin, P., Yang, Y., Jovicich, J., De Pisapia, N., Wang, X., Zuo, C.S., Levitt, J.J., 2016. Static and dynamic posterior cingulate cortex nodal topology of default mode network predicts attention task performance. Brain Imaging and Behavior 10, 212–225. https://doi.org/10.1007/s11682-015-9384-6

Liu, X., Duyn, J.H., 2013. Time-varying functional network information extracted from brief instances of spontaneous brain activity. Proceedings of the National Academy of Sciences 110, 4392–4397. https://doi.org/10.1073/pnas.1216856110

Mattar, M.G., Cole, M.W., Thompson-Schill, S.L., Bassett, D.S., 2015. A Functional Cartography of Cognitive Systems. PLOS Computational Biology 11, e1004533. https://doi.org/10.1371/journal.pcbi.1004533

Park, J.E., Jung, S.C., Ryu, K.H., Oh, J.Y., Kim, H.S., Choi, C.-G., Kim, S.J., Shim, W.H., 2017. Differences in dynamic and static functional connectivity between young and elderly healthy adults. Neuroradiology 59, 781–789. https://doi.org/10.1007/s00234-017-1875-2

Preti, M.G., Bolton, T.A., Van De Ville, D., 2017. The dynamic functional connectome: State-of-the-art and perspectives. NeuroImage 160, 41–54. https://doi.org/10.1016/j.neuroimage.2016.12.061

Rashid, B., Blanken, L.M.E., Muetzel, R.L., Miller, R., Damaraju, E., Arbabshirani, M.R., Erhardt, E.B., Verhulst, F.C., van der Lugt, A., Jaddoe, V.W.V., Tiemeier, H., White, T., Calhoun, V., 2018. Connectivity dynamics in typical development and its relationship to autistic traits and autism spectrum disorder. Human Brain Mapping 39, 3127–3142. https://doi.org/10.1002/hbm.24064

Rudie, J.D., Brown, J.A., Beck-Pancer, D., Hernandez, L.M., Dennis, E.L., Thompson, P.M., Bookheimer, S.Y., Dapretto, M., 2013. Altered functional and structural brain network organization in autism. NeuroImage: Clinical 2, 79–94. https://doi.org/10.1016/j.nicl.2012.11.006

Rudie, J.D., Shehzad, Z., Hernandez, L.M., Colich, N.L., Bookheimer, S.Y., Iacoboni, M., Dapretto, M., 2012. Reduced Functional Integration and Segregation of Distributed Neural Systems Underlying Social and Emotional Information Processing in Autism Spectrum Disorders. Cerebral Cortex 22, 1025–1037. https://doi.org/10.1093/cercor/bhr171

Schlesinger, K.J., Turner, B.O., Lopez, B.A., Miller, M.B., Carlson, J.M., 2017. Age-dependent changes in task-based modular organization of the human brain. NeuroImage 146, 741–762. https://doi.org/10.1016/j.neuroimage.2016.09.001

Song, J., Birn, R.M., Boly, M., Meier, T.B., Nair, V.A., Meyerand, M.E., Prabhakaran, V., 2014. Age-Related Reorganizational Changes in Modularity and Functional Connectivity of Human Brain Networks. Brain Connectivity 4, 662–676. https://doi.org/10.1089/brain.2014.0286

Sporns, O., 2013. Structure and function of complex brain networks. Dialogues Clin Neurosci 15, 247–262.

Tailby, C., Kowalczyk, M.A., Jackson, G.D., 2018. Cognitive impairment in epilepsy: the role of reduced network flexibility. Annals of Clinical and Translational Neurology 5, 29–40. https://doi.org/10.1002/acn3.503

Telesford, Q.K., Ashourvan, A., Wymbs, N.F., Grafton, S.T., Vettel, J.M., Bassett, D.S., 2017. Cohesive network reconfiguration accompanies extended training: Cohesive Network Reconfiguration. Human Brain Mapping 38, 4744–4759. https://doi.org/10.1002/hbm.23699

van den Heuvel, M.P., Hulshoff Pol, H.E., 2010. Exploring the brain network: A review on resting-state fMRI functional connectivity. European Neuropsychopharmacology 20, 519–534. https://doi.org/10.1016/j.euroneuro.2010.03.008

Watanabe, T., Rees, G., 2017. Brain network dynamics in high-functioning individuals with autism. Nature Communications 8, 16048. https://doi.org/10.1038/ncomms16048

Xu, Y., Lindquist, M.A., 2015. Dynamic connectivity detection: an algorithm for determining functional connectivity change points in fMRI data. Frontiers in Neuroscience 9. https://doi.org/10.3389/fnins.2015.00285

Yan, 2010. DPARSF: a MATLAB toolbox for “pipeline” data analysis of resting-state fMRI. Frontiers in System Neuroscience. https://doi.org/10.3389/fnsys.2010.00013

Ye, M., Yang, T., Qing, P., Lei, X., Qiu, J., Liu, G., 2015. Changes of Functional Brain Networks in Major Depressive Disorder: A Graph Theoretical Analysis of Resting-State fMRI. PLOS ONE 10, e0133775. https://doi.org/10.1371/journal.pone.0133775

Yerys, B.E., Gordon, E.M., Abrams, D.N., Satterthwaite, T.D., Weinblatt, R., Jankowski, K.F., Strang, J., Kenworthy, L., Gaillard, W.D., Vaidya, C.J., 2015. Default mode network segregation and social deficits in autism spectrum disorder: Evidence from non-medicated children. NeuroImage: Clinical 9, 223–232. https://doi.org/10.1016/j.nicl.2015.07.018

Zalesky, A., Fornito, A., Bullmore, E.T., 2010. Network-based statistic: Identifying differences in brain networks. NeuroImage 53, 1197–1207. https://doi.org/10.1016/j.neuroimage.2010.06.041

